# Multiple, not single, recipient muscle tendon transfers produce well-directed thumb-tip forces in lateral pinch grasp: a simulation study with application to restoration of improved grasp after tetraplegia

**DOI:** 10.1101/2025.01.20.633990

**Authors:** Oliver Garcia, Joseph D Towles

## Abstract

Thumb tendon transfer surgical procedures in patients with cervical spinal injury engage the paralyzed flexor pollicis longus (FPL) muscle to enable lateral pinch grasp. However, functional outcomes are mixed, in part because the FPL cannot produce force at the thumb tip to promote a stable grasp. We used simulation to investigate whether a multi-site tendon transfer, targeting sets of paralyzed muscles driven by a single donor muscle, could outperform a single-site tendon transfer with the FPL alone and restore lateral pinch. We formed 36 groups of 2 muscles, 84 groups of 3 muscles, and 126 groups of 4 muscles. We used nonlinear optimization and in-situ measurements of endpoint forces in 3 lateral pinch postures. We found that 116 of the 246 muscle groups outperformed the FPL for wide and narrow lateral pinch postures and a posture in between. On average, the 116 muscle groups produced endpoint forces 8° to 40° better directed than the FPL, and groups of four muscles statistically significantly outperformed the FPL’s force production. Our findings highlight the possibility of using multi-insertion site tendon transfers to restore grasp following cervical spinal cord injury.

## INTRODUCTION

Annually, up to 135,000 persons worldwide have a cervical spinal cord injury (B. B. Lee et al., 2014; *Spinal Cord Injury (SCI) Facts and Figures at a Glance*, n.d.). Resultant loss of hand function severely affects their quality of life. Tendon transfer surgical procedures restore lateral pinch grasp (Fox et al., 2018; Revol et al., 2002). In many such operations, the donor muscle is attached to the paralyzed flexor pollicis longus (FPL) muscle because it is the only thumb muscle that flexes all three thumb joints (the carpometacarpal joint, the metacarpophalangeal joint, and the interphalangeal joint), causes the least amount of radial-ulnar deviation (Paul Smutz et al., 1998), crosses the wrist, and is surgically straightforward to connect to (Fox et al., 2018; Revol et al., 2002). When the donor muscle is activated, the FPL transmits the donor muscle’s force, which influences the magnitude of the endpoint force during grasp contact. However, the direction of the endpoint force is determined by the musculoskeletal geometry (e.g., the ratio of muscle moment arms and the location of the muscle insertion point on the bone) of the FPL; therefore, the FPL plays a greater role than the donor muscle in determining the quality of grasp contact. Functional outcomes following lateral pinch restorative tendon transfers have been mixed, with persons after surgeries generating pinch forces from 1 to 2 N to 10 times as much (Johanson et al., 2016, p. 20; Waters et al., 1985). We think this is, in part, because the FPL’s musculoskeletal geometry does not give rise to a well-directed force at the thumb tip that would promote a slip-free, stable grasp (Towles, 2023; Towles et al., 2008). A well-directed endpoint force is closely aligned with the palmar force direction (Fig. 1). The FPL’s proximal force component is larger than its palmar force component; therefore, the FPL is more likely to promote slip in the proximal direction during grasp contact rather than promoting a pressing action that maintains grasp contact in the palmar direction (Fig. 1 illustrates the angular range resulting from FPL’s endpoint force components). This angular range suggests that when the donor muscle transmits its force through the FPL only, the resulting grasp force will be relatively weak.

**Figure 1:**
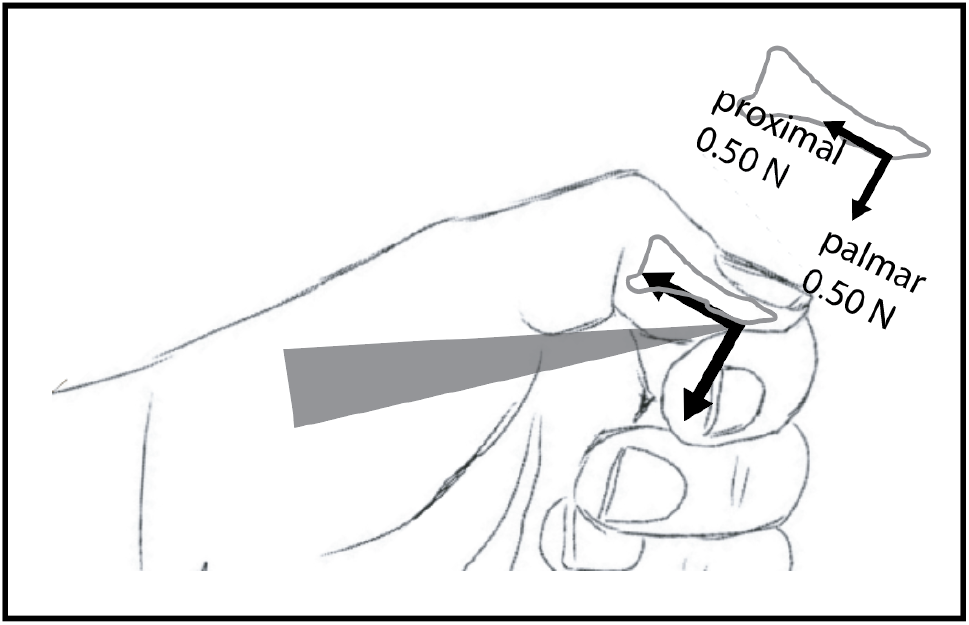
Angular Variation in Flexor Pollicis Longus (FPL) Endpoint Force When the Thumb Is Flexed. The gray triangle illustrates the 95% confidence interval around the FPL’s endpoint force direction, 46° to 55° relative to palmar direction. The height of the triangle represents the magnitude of the FPL’s endpoint force (2.8 N, Towles et al., 2008).

Multi-muscle control is required for strong and stable grasps (Johanson et al., 2001; Sharma & Venkadesan, 2022; Valero-Cuevas, 2000; Venkadesan & Valero-Cuevas, 2008). Thus, we think a small number of paralyzed muscles simultaneously driven by a donor muscle can produce a resultant endpoint force that is better directed than that of FPL alone during lateral pinch grasp, thus creating a stronger and more stable grasp. However, a general and long-standing tenet of tendon transfer procedures is to attach one donor muscle to one paralyzed recipient muscle (Hentz et al., 1983); that is, a single-site tendon transfer. We believe this tenet exists because of the difficulty of surgically planning the control of multiple recipient muscles by one donor muscle throughout the lateral pinch range of motion. Historically, computational musculoskeletal and cadaveric surgical simulation tools have not been used to plan lateral pinch tendon transfer procedures. We believe both simulations tools can be brought to bear to make multi-site tendon transfer surgical procedures (Fig. 2) tractable to solve.

**Figure 2:**
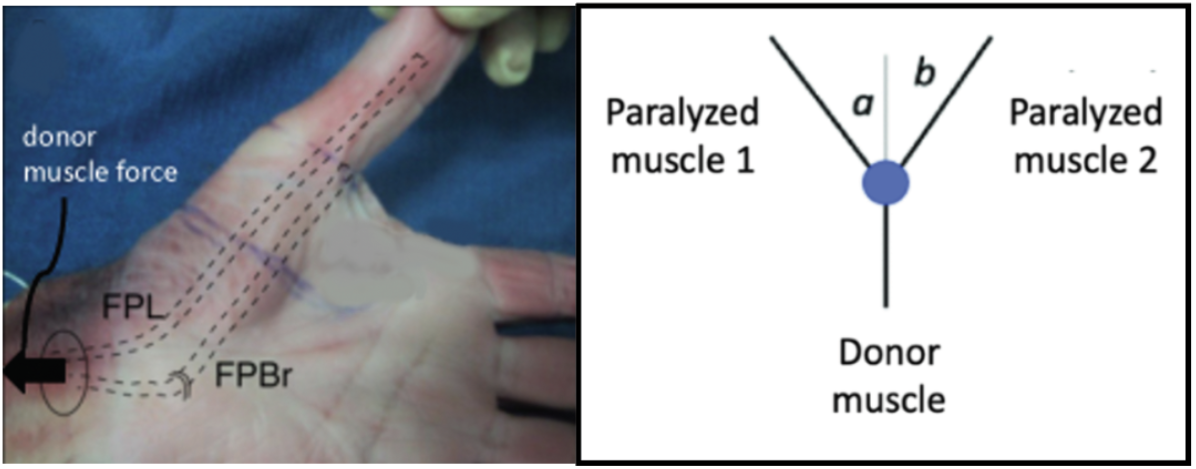
Multi-Site Tendon Transfer Illustration. In this arrangement, the donor muscle (not shown in the photo, left) is attached to two paralyzed recipient muscles—the flexor pollicis longus (FPL) muscle and the radial head of the flexor pollicis brevis (FPBr) muscle—and the donor muscle force is transmitted through the recipient muscles to actuate the thumb. The proportion of donor force transferred to each recipient muscle depends on the values of angles *a* and *b* (right). This figure was reproduced, in part, from Towles (Towles, 2023).

This study builds on our previous work (Towles, 2023) in which we investigated whether small groups of muscles could produce endpoint forces that were directed better than that of the FPL during lateral pinch grasp. In that simulation study, small groups of muscles, which represented sets of paralyzed recipient muscles, driven by a donor muscle produced endpoint forces better directed than that of the FPL when the thumb was in the flexed and mid-flexed, but not extended, postures. Functionally, each of the postures represents thumb positions for grasping small (flexed), mid-sized (mid-flexed), and large (extended) objects. Recipient muscles each transmitted the same level of donor muscle force (Towles, 2023). The goal of this current simulation study was to build on our previous study by allowing for the possibility that each paralyzed recipient muscle can transmit different levels of donor muscle force. We addressed two main questions: (1) Which of the tested muscle groups outperforms the FPL? and (2) To what extent do these muscle groups outperform the FPL?

## METHODS

### Data Collection Protocol

We simulated tendon transfer-driven lateral pinch force production using nonlinear optimization and in situ three-dimensional measurements of individual muscle endpoint forces in the flexed and extended thumb, as previously described (Towles, 2023; Towles et al., 2008).

Simulated tendon transfer procedures were novel in that, in each one, the donor muscle was attached to small groups of paralyzed muscles, representing a multi-site tendon transfer, rather than being attached to only one paralyzed muscle, representing a single-site tendon transfer. Tendon transfers procedures involved combinations of all nine thumb muscles: the extensor pollicis longus (EPL), the FPL, the extensor pollicis brevis (EPB), the radial (FPBr) and ulnar (FPBu) heads of the flexor pollicis brevis, the abductor pollicis brevis (APB), the adductor pollicis (ADP), the abductor pollicis longus (APL), and the opponens pollicis (OPP). From the nine muscles, we formed the maximum number of groups of 2 muscles, 36; of 3 muscles, 84; and of 4 muscles, 126. Thus, we simulated a multi-site tendon transfer surgery for each of the 246 muscle groups of paralyzed recipient muscles. Each simulated tendon transfer enabled lateral pinch force production in three representative thumb postures—flexed, extended, and mid-flexed—as proxies for endpoint force production in a narrow lateral pinch posture (flexed), wide lateral pinch posture (extended), and a posture in between (mid-flexed) throughout the flexion-extension plane (Fig. 3).

**Figure 3:**
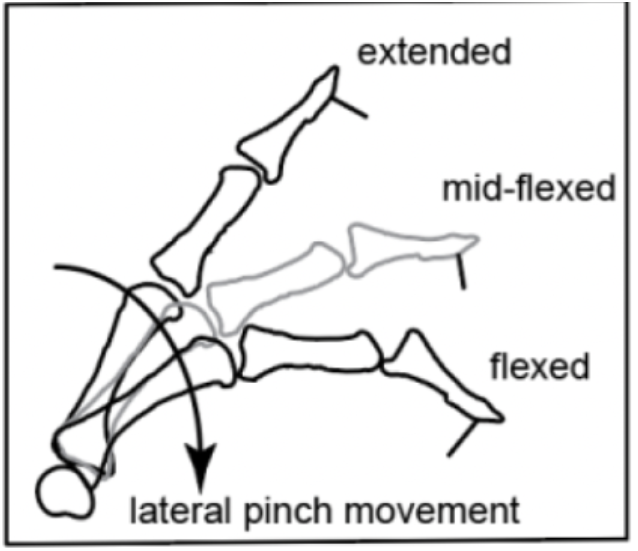
Flexed, Mid-Flexed, and Extended Thumb Postures. These postures span the lateral pinch or flexion-extension plane. The line segment normal to the distal phalanx is the palmer force direction, the most desirable direction in which to produce force for stable grasping. This figure was adapted from Towles (Towles, 2023).

As in a previous study (Valero-Cuevas et al., 2000), we used a cadaver-based mathematical model to model lateral pinch force production by the thumb in the postures of interest, expressed thus:

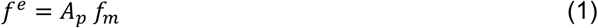

In EQ. 1, *A*_*p*_ is a *3×9* action matrix whose columns are the collection of the *3×1* median endpoint force vectors that each muscle produces when each muscle generates 1 N of force. That is, 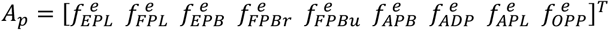 and *p* = 1, 2, or 3, signifying the flexed, mid-flexed, and extended thumb postures, respectively. *A*_*p*_ maps the *9×1* muscle force vector, 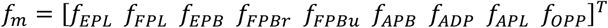, into the *3×1* resultant endpoint force vector, 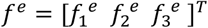, where directions 1, 2, and 3 correspond to the directions along the ulnar-radial, proximal-distal, and dorsal-palmar anatomical axes, respectively (Fig. 4). The ulnar, proximal, and dorsal directions are the positive anatomical directions in the model.

**Figure 4:**
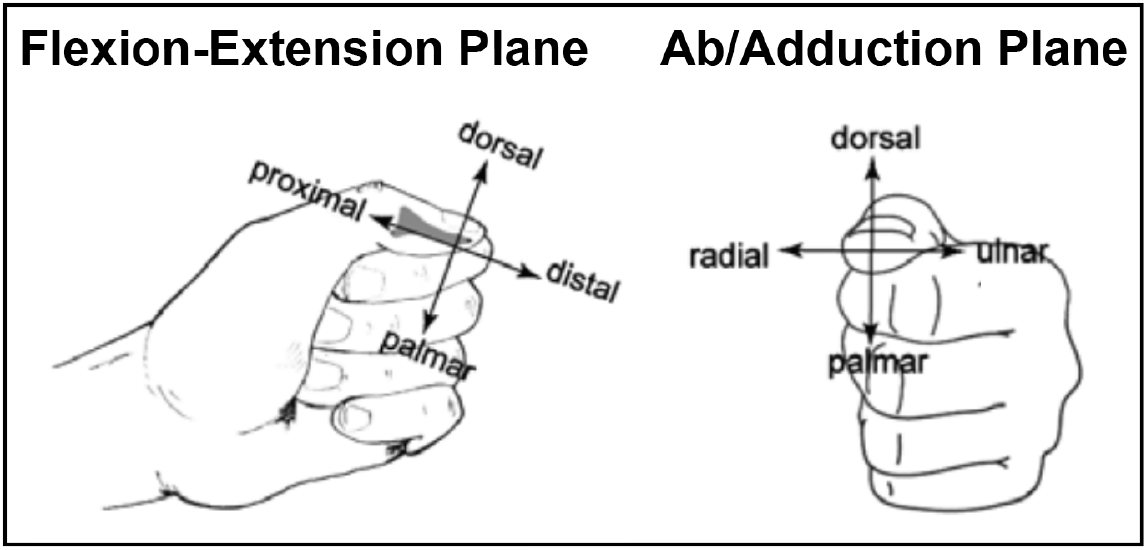
Thumb Tip Reference Frame. The three-dimensional reference frame is defined by the ulnar-radial, proximal-distal, and dorsal-palmar anatomical axes. This figure was adapted from Towles et al. (Towles et al., 2008)

The nonlinear optimization algorithm searched for *f*_*m*_ that minimized *θ*, the angle of the endpoint force that a muscle group produces relative to the palmar force direction. *θ* was given by

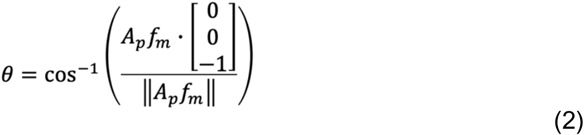

In EQ. 2, muscle forces, *f*_*m,i*_, ranged from zero to their maximum isometric force and were determined by the product of maximum tissue stress (35 N/cm^2^, (Zajac, 1989)) and each muscle’s physiological cross-sectional area (Brand et al., 1981; Jacobson et al., 1992; Lieber et al., 1992). For a muscle to be considered a contributor to the resultant endpoint force of a muscle group, the muscle had to produce at least 0.1 N of force.

For each muscle group in the flexed posture, if *θ* was less than the lower limit of the 95% confidence interval (CI) around the FPL’s endpoint force direction (i.e., less than 46°), then that muscle group, its generated endpoint force (using EQ. 1), and muscle forces were recorded. Similarly, for the mid-flexed and extended postures, if *θ* in each case was less than the lower bounds of the 95% CIs around the FPL’s endpoint force directions in those postures (i.e., less than 59° and 72°, respectively), then those muscle groups, their generated endpoint forces (using EQ. 1), and muscle forces were recorded.

### Data Analysis

To determine whether a muscle group produced an endpoint force directed more toward the palmar direction than that of the FPL in each posture, we compared each muscle group’s endpoint force direction to the 95% CI around the FPL’s endpoint force direction in each of the three postures.

In each posture, if multiple muscle groups outperformed the FPL, we used a one-way analysis of variance (ANOVA, α = 0.05) to compare endpoint force directional improvements among muscle groups. For statistically significant directional differences, Tukey post hoc tests were used to determine which inter-group comparisons were statistically significant.

To understand the contribution of individual muscles on force directional improvements in each posture, the mean angular improvement in groups with each muscle was compared to the mean angular improvement in the same groups without each of those muscles. Student t tests with a Bonferroni correction for each of the nine thumb muscles determined whether including a muscle in a group was associated with a significant difference in angular improvement, with α = 0.05/9 = 0.006.

FPBr and ADP were the only muscles whose inclusion was associated with a significant improvement in mean endpoint force angle when present in a muscle group (Table 2). On average, groups with ADP were 5.8° closer to the palmar direction than those that did not include ADP. Groups with FPBr improved by a mean of 5.37°.

## RESULTS

### Overall Force Angle Improvement

We found 116 (47.2%) of the 246 muscle groups examined had better-directed endpoint forces than the FPL, with directions less than 46°, 59°, and 72° in the flexed, mid-flexed, and extended postures, respectively. When averaged across the three postures, group endpoint forces ranged from 17° to 49° relative to the palmar direction (Fig. 5, Table 1), an mean improvement of 8° to 40° over the FPL’s mean endpoint force direction across all three postures. Several muscle groups generated endpoint forces exactly in the palmar direction (0°) in the flexed and mid-flexed postures. These muscle groups are the last 28 groups boxed in Table 1. In the extended posture, no muscle groups generated endpoint forces less than 53° relative to the palmar direction.

**Figure 5:**
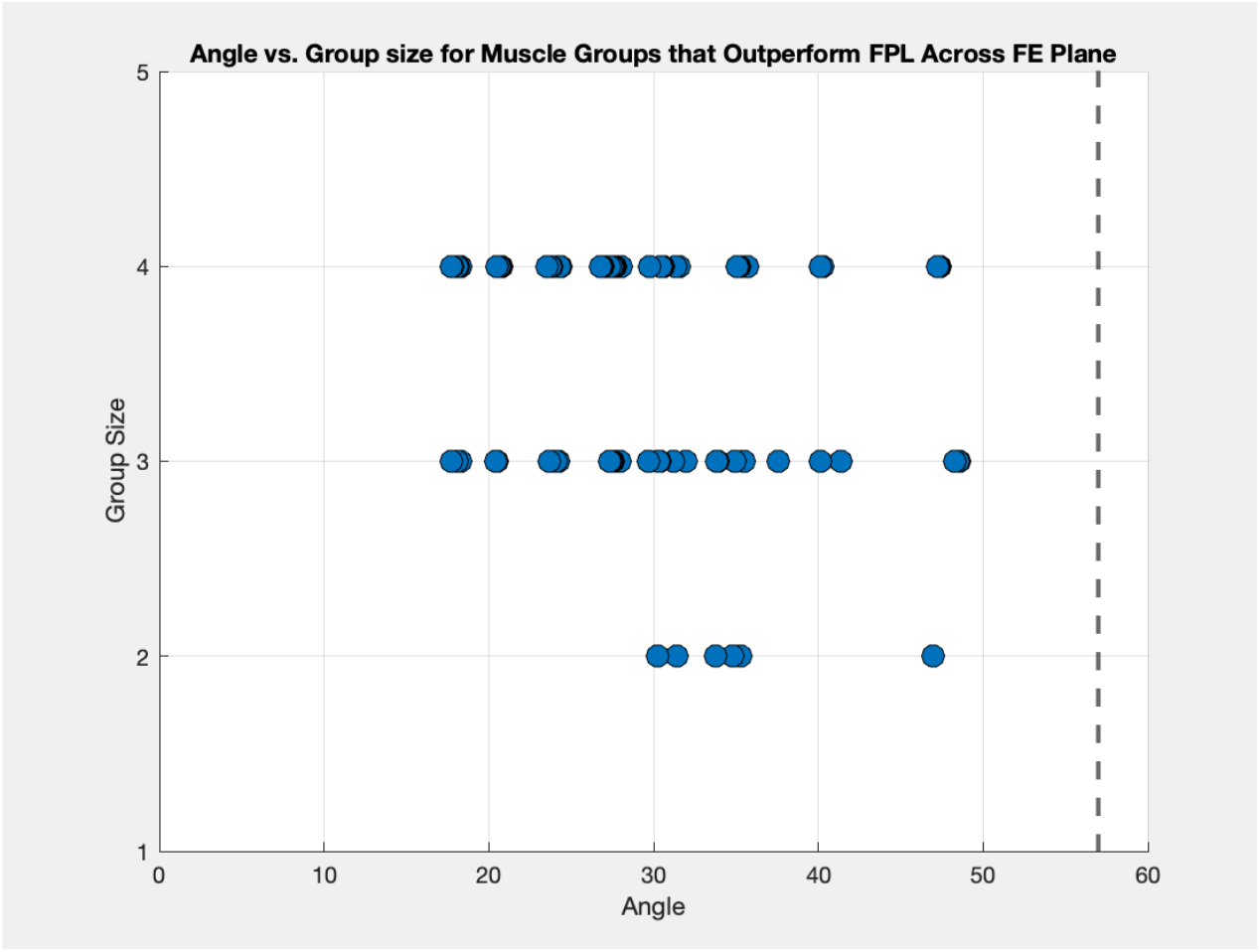
Muscle Group Endpoint Force Angles. Each muscle group that outperformed the flexor pollicis longus (FPL) muscle in all 3 postures is presented with the angle of the force generated by that group (averaged across all postures) and the number of muscles involved in each group. The FPL’s mean endpoint force angle, when activated alone, is shown as the vertical line at 57°.

**Table 1:**
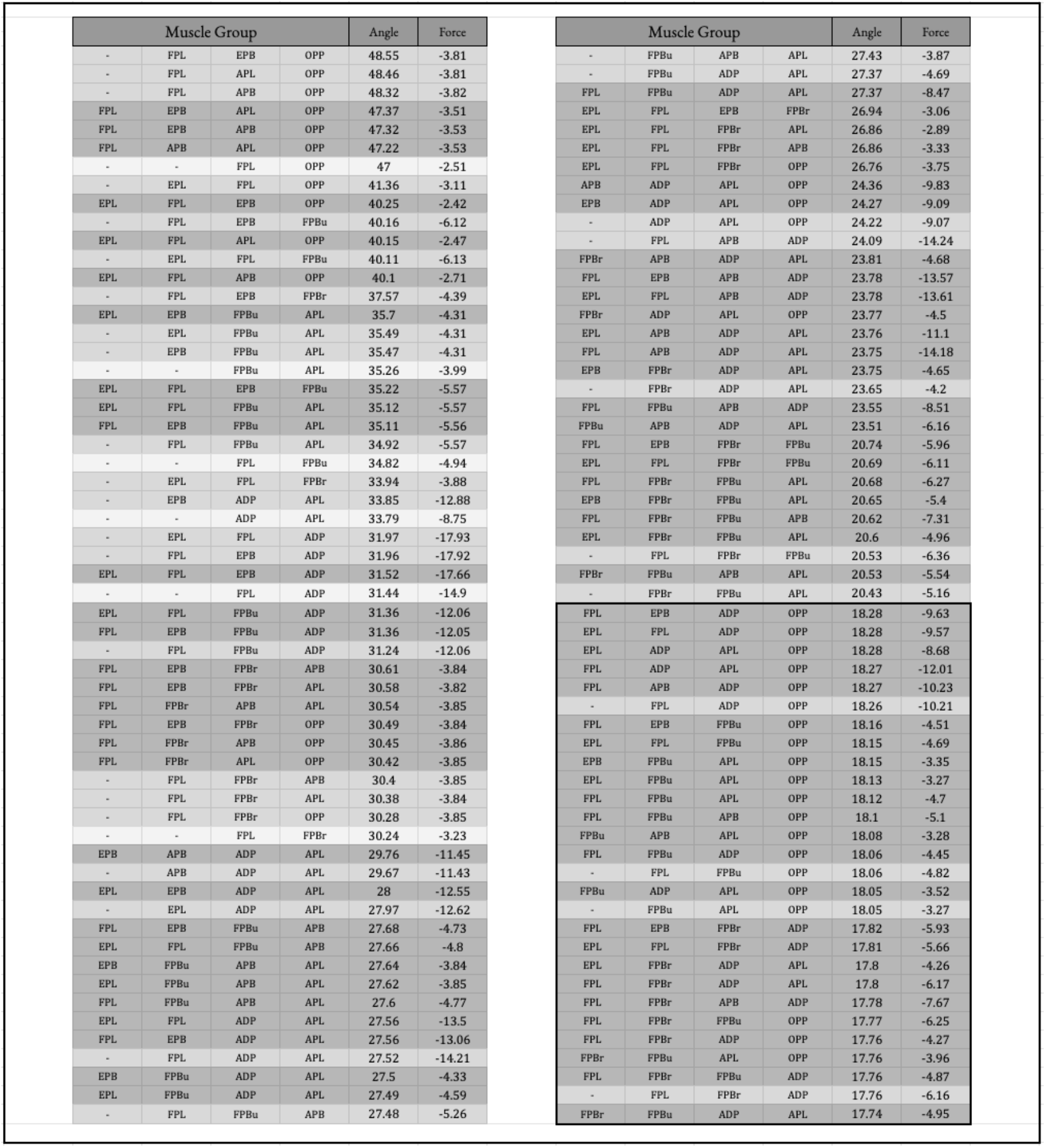
Endpoint Force Directions of the 116 Muscle Groups that Outperformed the Flexor Pollicis Longus (FPL) Muscle in All Three Postures. The angle of the endpoint force vectors generated in each posture were averaged across all postures for each group. The FPL produced an average force directed 57° from the palmar direction.

### Angular Improvement Among Muscle Groups

Of the 116 muscle groups that outperformed the FPL, there were 6 groups of 2 muscles, 33 groups of 3 muscles, and 77 groups of 4 muscles. Across all postures, the groups of 2 muscles, as a whole, produced endpoint forces directed at 35.4° with respect to the palmar direction; the groups of 3 muscles, 30.8°; and the groups of 4 muscles, 25.5°. The across-posture mean of the groups of 4 muscles was less than that of the groups of 3 muscles (*p* = 0.004) and 2 muscles (*p* = 0.009) (Fig. 6).

**Figure 6:**
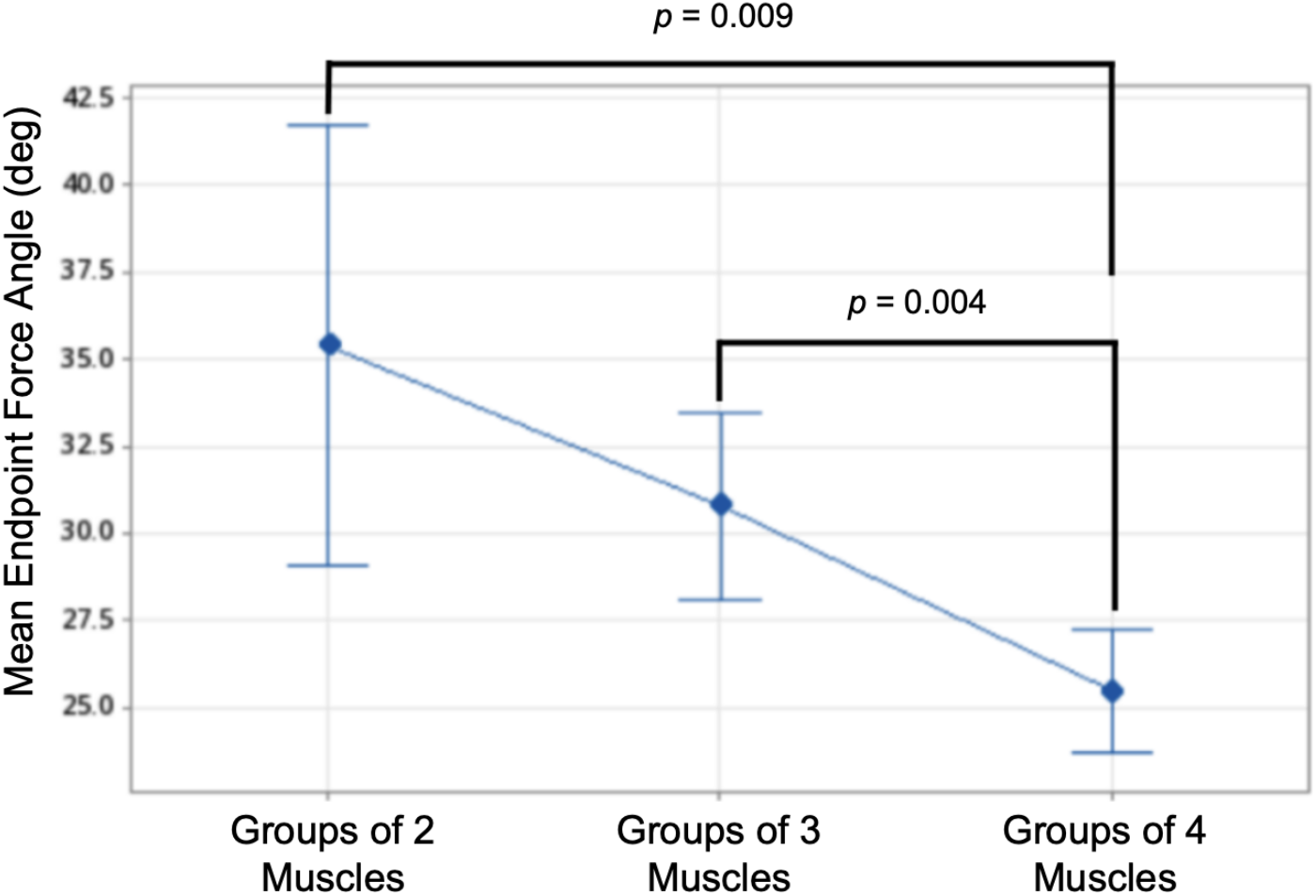
Mean Endpoint Force Angles by Group Size. The graph presents 95% confidence intervals.

### Angular Improvement by Muscle Inclusion

Individual muscles did not appear an equal number of times in the muscle groups that outperformed the FPL across all three postures. That is, the FPL appeared 78 times; APL, 60; FPBu, 51; ADP, 51; FPBr, 40; OPP, 40; EPL, 33; EPB, 33; and APB, 33. Groups containing FPBr and ADP had the lowest mean endpoint force angles, 23.98° and 24.23°, respectively (Table 2). FPBr and ADP were the only muscles whose inclusion was associated with a significant improvement in mean endpoint force angle when present in a muscle group (Table 2). On average, groups with ADP were 5.8° closer to the palmar direction than those that did not include ADP. Groups with FPBr improved by a mean of 5.37°.

**Table 2:** Muscles With Significant Impact on Force Angle. The average endpoint force angles of groups that contained each muscle compared with the average angle of groups from which each muscle was absent. The inclusion of adductor pollicis (ADP) and the radial head of the flexor pollicis brevis (FPBr) in a muscle group is each associated with a significantly smaller force angle.

| Muscle | No. Inclusions | Mean Angle When Present | Mean Angle When Absent | P Value |
| --- | --- | --- | --- | --- |
| ADP | 51 | 24.24 | 30.06 | 0.001 |
| FPBr | 40 | 23.98 | 29.35 | 0.001 |

## DISCUSSION

This study builds on our previous investigations on how to improve tendon transfer surgical procedures aimed at restoring lateral pinch grasp in persons with tetraplegia (Towles, 2023; Towles et al., 2008). The current approach, a tendon transfer procedure that engages the paralyzed FPL to actuate the thumb, can lead to unstable grasp contact and weak pinch forces. Our group’s previous work and this study aimed to address this challenge in two ways, using simulation. First, we reimagined grasp-restorative tendon transfer surgeries by using one donor muscle to engage multiple paralyzed recipient muscles that actuate the thumb. We term this type of operation a “multi-site tendon transfer” because the donor muscle engages multiple paralyzed muscles. Second, we considered thumb tip movement and force production throughout the entire plane of lateral pinch motion to optimize tendon transfers for both planar movement of the thumb and stable grasp contact. To the best of our knowledge, designing grasp-restorative tendon transfer procedure through the lens of the endpoint mechanical behavior of the thumb is new and aligns well with concepts in robotics well-suited to analyze grasp motion, contact force production, and stability (Lynch & Park, 2017). Toward this end, the primary goal of this study was to explore whether small groups of muscles produced endpoint forces that were directed better than that of the FPL when the thumb was positioned in flexed, mid-flexed and extended postures and when forces transmitted by recipient muscles were allowed to vary from posture to posture. Muscle endpoint force production in the mid-flexed posture was computed to be the mean of the muscle endpoint forces produced in the flexed and extended postures.

In this study, we addressed two main questions: (1) Which of the tested muscle groups outperform the FPL? and (2) To what extent do these muscle groups outperform the FPL? Of the 246 muscle groups tested across the three postures, we found 116 groups produced endpoint forces directed more toward the palmar direction than the FPL’s endpoint force (Fig. 5, Table 1). When averaged across the three postures, group endpoint forces ranged from 17° to 49° relative to the palmar direction (Fig. 5). This represented an mean improvement of 8° to 40° over the endpoint force direction that the FPL alone generated when averaged across all three postures.

Every muscle contributed to at least 33 successful muscle groups, with the FPL contributing to the most number of groups, 78. Every muscle contributed to multiple successful muscle groups likely because the set of nine individual thumb muscle endpoint forces spans the positive and negative directions of each of the three anatomical endpoint axes (ulnar-radial, proximal-distal, dorsal-palmar) of the endpoint force workspace (Towles et al., 2008). This characteristic enabled many different muscle combinations to outperform the FPL. In practice, the simultaneous action of multiple muscles enables well-directed endpoint forces (Johanson et al., 2001; Sharma & Venkadesan, 2022; Valero-Cuevas, 2000; Venkadesan & Valero-Cuevas, 2008). Given the FPL appeared in 78 of 116 successful muscle groups, the endpoint forces of the other muscles of those groups were coordinated with that of the FPL in such a way as to compensate for the FPL’s proximal force component and to complement the FPL’s palmar force component. In doing so, muscle groups compensated for the FPL’s tendency for unstable grasp contact (proximal force component) and complemented the FPL’s tendency for stable grasp contact (palmar force component). The FPBr and the ADP were the only two muscles whose inclusion was, on average, associated with a significant improvement in mean endpoint force angle, suggesting that these muscles most constructively combine with the FPL. The finding that these muscles are intrinsic agrees with our earlier simulation work that showed the beneficial endpoint force production characteristics of intrinsic muscles in the design of grasp-restorative procedures to improve grasp in persons following tetraplegia (Towles et al., 2008).

More groups of four muscles than those of two and three muscles outperformed the FPL and also did so with a resultant endpoint force direction more closely aligned with the palmar force direction (Fig. 6). This was the case because the groups of four muscles consisted of all of the successful three-muscle groups plus an additional helpful muscle. Generally speaking, muscle endpoint force production by a larger group of muscles can create a more well-directed endpoint force than can smaller groups of muscles because of the wider pool of muscle endpoint force vectors available. This idea is generally supported by the understanding that the actions of multiple muscles enable well-directed endpoint forces and that the capacity for directional improvement increases as the number of muscles increases. This understanding also explains why groups of four muscles outperformed groups of two muscles.

The finding that some muscle groups achieved muscle endpoint forces that aligned perfectly with the palmar direction in the flexed and mid-flexed postures (the last 28 groups boxed in Table 1) as compared with no such muscle groups in the extended posture is likely indicative of the extent to which the set of individual muscle endpoint forces in the flexed and extended thumb span the three-dimensional endpoint force space. In the flexed posture, thumb muscles produced force in all endpoint force directions with no direction experiencing a disproportionate amount of endpoint force magnitude (Towles et al., 2008). However, in the extended posture, thumb muscles did not produce force in all directions and did produce force in disproportionate amounts in other directions (Towles, 2023). Specifically, no muscle produced force in the distal direction, two of nine muscles produced small endpoint force in the dorsal direction compared with the other directions, and most muscles produced an inordinate amount of force in the proximal direction relative to other directions (Towles, 2023). These characteristics make it likely that muscle combinations in the flexed and mid-flexed postures, and not in the extended posture, could produce endpoint force exactly in the palmar direction.

As far as we know, this simulation study is the first to apply an endpoint force-based biomechanics approach to designing tendon transfer operations aimed at restoring lateral pinch grasp throughout the flexion-extension plane following cervical spinal cord injury. This study highlights the possibility of using multi-site tendon transfers to restore the ability to grasp. Investigating the surgical feasibility of engaging multiple recipient muscles, including intrinsic muscles (S. K. Lee & Wisser, 2012), in a tendon transfer operation and determining how to implement the appropriate coordination of donor muscle force across recipient muscles are the future steps to consider. As to the latter, angular changes should be controlled or predicted (Fig. 2) to appropriately modulate the transmission of donor muscle force to recipient muscles.

## ACKOWLEDGEMENT

Funding for this project was provided by the Provost’s Office at Swarthmore College (Garcia)

